# Enhanced detection of low-expressed miRNAs in *Leishmania*-infected macrophages through RNA fractionation and RT-qPCR optimization

**DOI:** 10.1101/2025.08.20.671241

**Authors:** S. Maestrini, H. Lecoeur, C. Gharsallah, A. Diotallevi, G. F. Späth, L. Galluzzi, E. Prina

## Abstract

MicroRNAs (miRNAs) play critical roles in regulating host responses to *Leishmania* infections, yet accurate detection of low-abundance miRNAs remains challenging. This study evaluated the impact of RNA fractionation and RT-qPCR protocol optimization on miRNA quantification in *Leishmania amazonensis*-infected murine macrophages. Using a panel of nine infection-associated miRNAs, we compared small RNA and total RNA fractions for their ability to detect weakly expressed targets. Small RNA consistently provided greater sensitivity and specificity, particularly when combined with a modified RT-qPCR protocol. These findings underscore the importance of RNA preparation methods for studying miRNA dynamics in infectious disease contexts and support improved approaches for detecting biologically relevant, low-expressed miRNAs.

## Introduction

MicroRNAs (miRNAs) are small non-coding RNAs that serve as pivotal post-transcriptional regulators of the host response, orchestrating gene expression in response to a myriad of physiological and pathological stimuli, including parasitic infections. Among these, obligate intracellular protozoan parasites of the genus *Leishmania* have evolved sophisticated mechanisms to manipulate host miRNA profiles, thereby modulating immune responses to favor their survival and proliferation within mammalian macrophages. Recently, epigenetic subversion such as DNA methylation, histone modification and the modulation of host miRNAs have been identified as a strategy employed by *Leishmania* to evade host defenses (Marr et al., 2014, Diotallevi et al., 2018, Lecoeur et al., 2020, Gutierrez Sanchez et al., 2023; Gharsallah et al., 2024). Cutaneous leishmaniasis (CL) is the most common form of *Leishmania* infection, typically presenting as localized skin lesions. These lesions can become chronic, resulting in significant tissue destruction and disfigurement. In CL, lesions may heal spontaneously with tissue fibrosis or progress to disseminated cutaneous leishmaniasis (DCL) or mucocutaneous leishmaniasis (ML). *Leishmania amazonensis* is one of the causative species of CL and DCL, characterized primarily by numerous heavily infected macrophages. The aim of this study was to determine the most appropriate RNA fraction—small RNA or total RNA—for the accurate analysis of low-abundance microRNAs using reverse transcription quantitative PCR (RT-qPCR) and to optimize the associated experimental conditions. The technical approach includes two key components: (i) miRNA isolation using affinity column-based kits for small and large RNA and magnetic-bead purification for total RNA extraction, and (ii) optimization of RT-qPCR for the sensitive detection of miRNAs, including those with very low expression levels. A panel of nine microRNA targets was selected for the study based on prior RNA-seq data from Gharsallah et al. (2024), which focused on identifying and characterizing microRNAs involved in the macrophage response to *L. amazonensis* infection in vitro.

## Materials and methods

### Ethics statement

Work on animals were conducted in compliance with French and European regulations on care and protection of laboratory animals (EC Directive 2010/63, French Law 2013-118, February 6th, 2013). All animal experiments were approved by the Ethics Committee and the Animal welfare body of Institut Pasteur and by the “Ministère de l’Enseignement Supérieur, de la Recherche et de l’Innovation” (project n°#49234). Bone marrow-derived macrophages (BMDMs) were obtained from healthy mice and infected macrophages were isolated from lesions of experimentally infected animals, ensuring minimal animal suffering and adherence to ethical standards for biomedical research.

### Production of bone-marrow derived macrophages

Bone marrow cells were isolated from the tibias and femurs of female C57Bl/6JRj mice and cultured in complete DMEM medium with high glucose concentration and supplemented with glucose, pyruvate, L-Alanyl-L-Glutamine, 10% heat-inactivated fetal calf serum, antibiotics, and β-mercaptoethanol, as described by Lecoeur et al., 2022. Cells (1 × 10^6^/mL) were differentiated into macrophages in the presence of 50 ng/mL mouse recombinant CSF1 and incubated in bacteriological Petri dishes at 37°C under 7.5% CO_2_. After 6 days, adherent BMDMs were detached using PBS with 25 mM EDTA, collected, and resuspended in complete medium supplemented with 30 ng/mL rm-CSF1. BMDMs were then seeded in 100 mm tissue culture dishes for RNA isolation.

### Parasite isolation and macrophage infection

mCherry-transgenic *L. amazonensis* amastigotes (strain LV79) were isolated from infected footpads of Swiss nude mice as described in Lecoeur et al., 2020. On day 6 of differentiation, BMDMs were seeded and allowed to adhere for 5 hours at 37°C. The macrophages were then infected with lesion-derived *L. amazonensis* amastigotes at a multiplicity of infection (MOI) of 4. Following infection, macrophages were cultured at 34°C and classified as Infected (I) BMDMs and Uninfected (UI) BMDMs.

### RNA extraction and RNA quality evaluation

Fractions of large (> 200 nucleotides) and small (< 200 nucleotides) RNAs were isolated from BMDMs at 24, 48 and 72 hours post infection (PI) using the NucleoSpin miRNA kit (Macherey-Nagel) while total RNAs were extracted using the MagMax mirVana Total RNA Isolation Kit (Applied Biosystems), following either the manufacturer’s protocol or modified in-house procedures. RNA quality was assessed using optical density measurements on a Nanodrop device (Kisker, http://www.kisker-biotech.com). Further quality assessment of selected RNA fractions was performed using the Agilent RNA 6000 Nano Total RNA Kit and the Agilent Small RNA Kit on a Bioanalyzer 2100 (Agilent) at the Biomics platform of Institut Pasteur. The analyses were conducting according to the manufacturer’s protocols.

### MicroRNA expression analysis using RT-qPCR

Nine microRNAs were analyzed in small RNA, large RNA and total RNA samples. cDNA synthesis was performed from all samples using the *mirCURY LNA miRNA SYBR Green PCR* kit (Cat. No. / ID: 339320), which enables the analysis of multiple miRNAs from a single reverse transcribed sample. Initially, 10 ng of RNA was used for a 10 µL reverse transcription (RT) reaction according to the manufacturer’s protocol. Subsequently, a 1:60 dilution of the resulting cDNA was employed for target detection by qPCR (referred to as Classical protocol). In a second approach, in-house modifications were introduced — increasing the RNA input for RT to 130 ng and reducing also the cDNA dilution for qPCR to 1:6 for low expressed miRNA — in order to enhance assay sensitivity (referred to as Protocol 1 and 2, respectively). The different protocols are summarized in Table 1. For the low abundance target miRNA-686, a synthetic positive control (mmu-miR-686 miRCURY LNA miRNA PCR assay, ref YP00205454, Qiagen) was used. The synthetic miRNA mimic was resuspended according to the Manufacturer’s instructions and 6.7 pg was used in RT reaction. Additionally, the UniSp6 synthetic molecule, which is not found naturally in biological samples, was used as a spike-in control to monitor the efficiency of the reverse transcription reaction and the quality of the resulting cDNA. The targets selected for this study were based on previous RNAseq experiments and their mean read counts, reported in Table 2. The following miRNAs were analyzed: mmu-miR-686, mmu-miR-16-5p, mmu-let-7f-5p, mmu-miR-103-3p, mmu-miR-99b-5p, mmu-miR-342-3p, mmu-miR-26a-5p, mmu-miR-674-5p and mmu-miR-7b-5p. Each qPCR reaction was performed in technical duplicates, with a final volume of 10 µl per replicate. Amplifications and melting curves were carried out on the LightCycler 480 system using the thermal cycling conditions recommended by the kit manufacturer. Quantitative cycle (Cq) values and melting curves for each sample from each target were compared and analyzed to identify the most suitable starting material and the optimal conditions for miRNA analysis.

**Table 1.** Improvements of the classical protocol of miRNA analysis: The classical protocol uses 10 ng RNA with a 1:60 cDNA dilution. Protocol 1 increases the RNA input to 130 ng while Protocol 2 combines this higher RNA input with a reduced cDNA dilution of 1:6 to enhance the detection of low-abundance miRNAs. In each protocol the amount of cDNA used in the qPCR reaction was 3:10 (cDNA:total reaction volume).

|  | Amount of RNA for cDNA synthesis | cDNA dilution for qPCR reactions |
| --- | --- | --- |
| **Classical protocol** | 10 ng | 1:60 |
| **Protocol 1** | 130 ng | 1:60 |
| **Protocol 2** | 130 ng | 1:6 |

**Table 2.** Transcriptomic read counts of selected microRNAs. The table shows mean read counts of selected miRNAs in three *L. amazonensis* infected (I) and uninfected (UI) biological replicates of BMDMs from previous RNAseq experiments described in Gharsallah C. et al., 2024. Fold change (FC) represents the ratio between I and UI reads, while log2FoldChange (log2FC) indicates the logarithmic transformation of this ratio. Statistical significance was assessed using the Benjamini-Hochberg method to calculate adjusted p-values (padj), with values below 0.05 considered indicative of differential expression.

| Id | UI reads | I reads | FC | log2FoldChange | padj |
| --- | --- | --- | --- | --- | --- |
| **mmu-miR-342-3p** | **55,522** | **31,602** | **0.593** | **-0.753** | **0.0060** |
| **mmu-miR-99b-5p** | **4,508** | **1,926** | **0.42** | **-1.252** | **1.75E-10** |
| **mmu-miR-26a-5p** | **28,564** | **22,269** | **0.78** | **-0.359** | **0.0028** |
| **mmu-miR-674-5p** | **2,312** | **3,438** | **1.476** | **0.562** | **0.2697** |
| **mmu-miR-7b-5p** | **16** | **15** | **0.897** | **-0.157** | **1** |
| **mmu-let-7f-5p** | **38,657** | **48,667** | **1.258** | **0.331** | **0.1267** |
| **mmu-miR-103-3p** | **20,125** | **15,632** | **0.77** | **-0.377** | **0.4956** |
| **mmu-miR-16-5p** | **329,632** | **318,046** | **0.965** | **-0.051** | **1** |
| **mmu-miR-686** | **3** | **3,953** | **1206.017** | **10.236** | **4.21E-52** |

## Results and Discussion

### Optimization of RT-qPCR conditions enhances detection of low-abundance miR-686 in uninfected macrophages

Nine miRNA targets were analyzed by RT-qPCR in order to determine the most suitable fraction between small and total RNAs for detecting low-abundance targets. Among them, mmu-miR-686 was identified as a low-expressed miRNA in uninfected (UI) BMDMs, but up-regulated in *L. amazonensis* infected (I) BMDMs, as described in Gharsallah C. et al., 2024. Transcriptomic data showed an over 1,000-fold increase in abundance (from 3 normalized reads in UI BMDMs to 3,953 in I BMDMs) (Table 2). To improve the detection of this target in UI BMDMs, the classical protocol was optimized in two ways: Protocol 1 increased the RNA input during cDNA synthesis to 130 ng and Protocol 2 further improved sensitivity by reducing the cDNA used in the PCR reaction to 1:6. Amplification curves show that both the classical protocol and Protocol 1 fail to generate detectable signals below the threshold of 40 cycles, indicating poor sensitivity for low-abundance targets (Figure 1). In contrast, the optimized Protocol 2 shifts Cq values to earlier cycles (Cq < 40), demonstrating improved detection efficiency.

**Figure 1.**
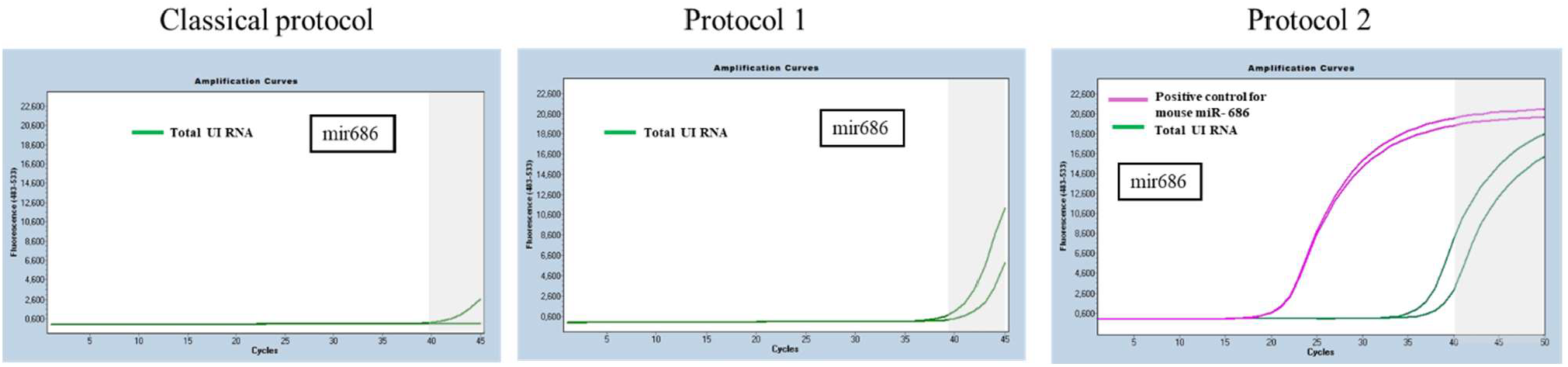
Improved detection of miR-686 in uninfected total RNAs samples. The classical protocol was optimized by increasing RNA input for cDNA synthesis (Protocol 1) and reducing cDNA dilution in qPCR (Protocol 2) to better detect miR-686 in uninfected (UI) BMDMs. The figure represents two technical replicates of UI samples, and the positive control for mouse miR-686. The grey area in the amplification plots marks cycles beyond 40, below the detection threshold. While the classical protocol and Protocol 1 show no signal, Protocol 2 shifts amplification before cycle 40 in both technical replicates, indicating improved sensitivity.

### Use of small RNA fraction eliminates non-specific amplification of low-abundance miR-686

Interestingly, the amplicon obtained from the total RNA fraction showed two distinct peaks in melting curve analysis (Figure 2B1), suggesting the presence of non-specific products. To address this, the same protocols were applied using the small RNA fraction. Under the conditions of Protocol 2, as described in the Methods section, the small RNA fraction proved more effective in reducing non-specific products for this low-expressed target (Figure 2B2).

**Figure 2.**
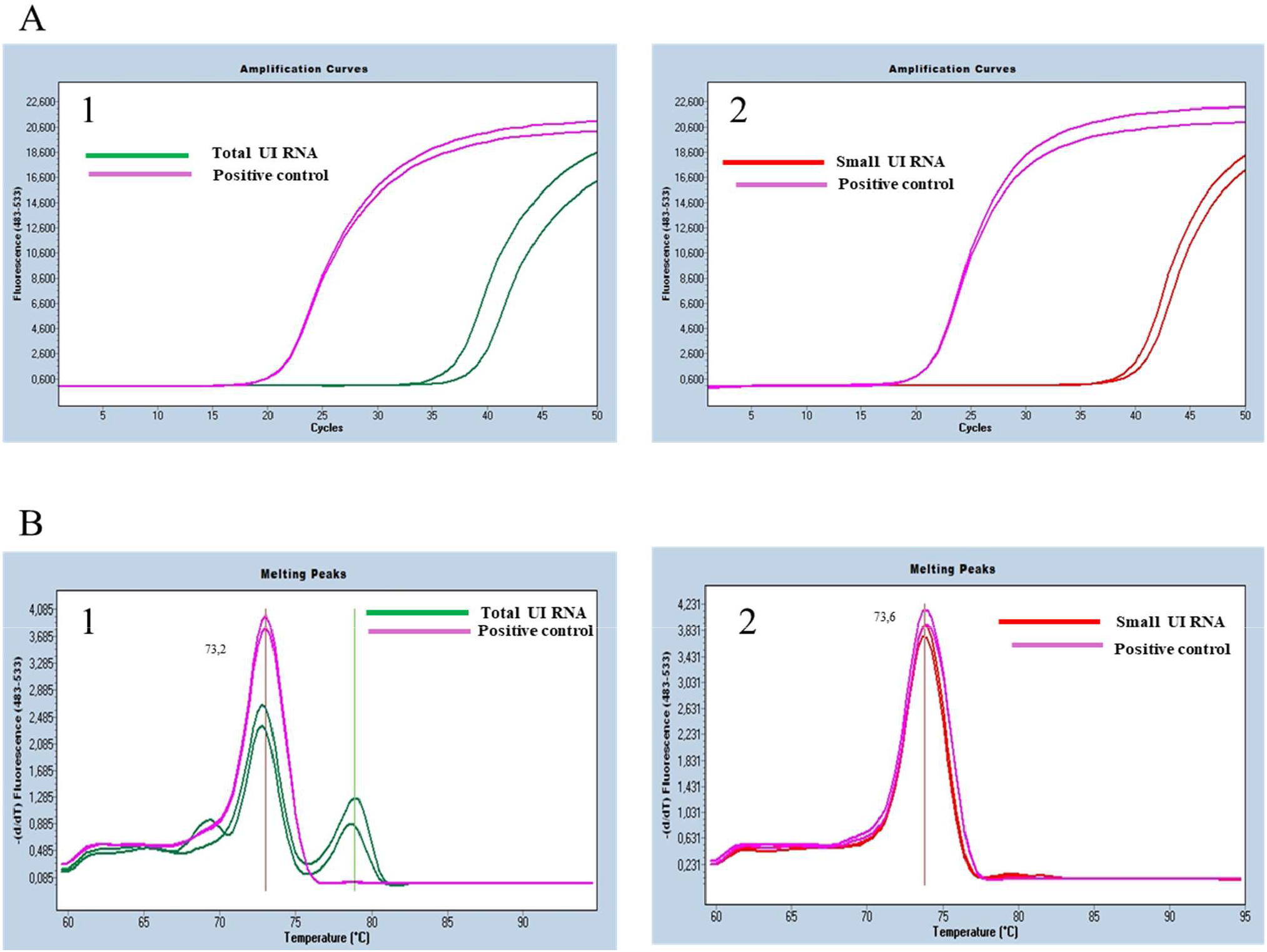
Comparison of amplification and melting curves for miR-686 in small and total RNAs samples from uninfected BMDMs. Amplification (A) and melting (B) curves for miR-686 using Protocol 2 in Total RNA (green) and Small RNA (red) fractions, compared to positive controls (purple). Panels A1 and B1 show data from Total RNA samples, whereas panels A2 and B2 correspond to small RNA samples. The specific miR-686 amplicon was detected in uninfected (UI) BMDMs only when using the small RNA fraction. Data represent two technical replicates of UI samples along with the positive control for mouse miR-686.

### Specific amplification of low-abundance miRNAs is achieved only with small RNA fraction

Regarding the specificity of the amplification products obtained with the other targets under investigation, both RNA fractions produced specific products for highly expressed target, as indicated by the presence of a single melting curve in panels A3 and B3 of Figure 3. However, for low-expressed targets, non-specific melting curves were observed in the total RNA fraction (panels A1 and A2 of Figure 3), as well as in the large RNA fraction that was performed in parallel (panels A and B of Figure S2).

**Figure 3.**
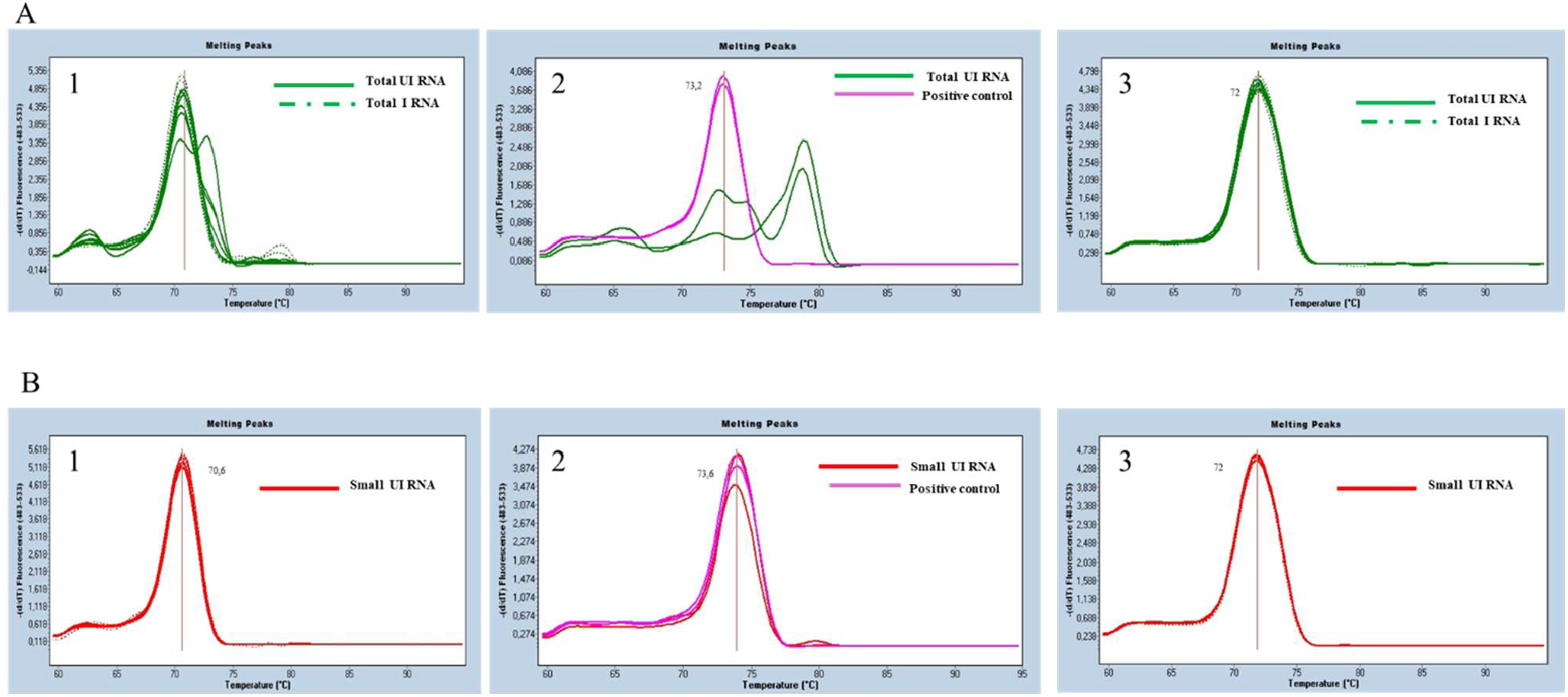
Comparison of melting curves from total and small RNA fractions in *L. amazonensis*-infected and uninfected BMDMs for both high- and low-abundance miRNAs. A1 and B1 panels show melting curves of miR-7b-5p from total and small RNA fractions, respectively, in both uninfected (UI) and infected (I) samples at 24 h, 48 h, and 72 h PI. A2 and B2 panels show melting curves of miR-686 in total and small RNA fractions from UI samples at 72 h PI. A3 and B3 panels show melting curves of miR-26b in total and small RNA fractions from UI and I samples at 24 h, 48 h and 72 h PI. While both RNA fractions yielded specific amplification for highly expressed targets (panels 3), only the small RNA fraction produced specific melting peaks for low-abundance miRNAs such as miR-7b-5p and miR-686 (panels B1 and B2) in UI macrophages, whereas total RNA showed non-specific profiles (panels A1 and A2).

### Small RNA fraction yields lower Cq values across targets

For all nine targets analyzed, RT-qPCR using small RNAs as the starting material consistently yielded lower Cq values compared to both the total RNA fraction (Figure 4) and the large RNA fraction (Figure S1). This trend was observed for both highly and lowly expressed targets, with the exception of miR-686. For this miRNA, Cq values for the small and total RNA fractions were comparable. However, as previously shown, only the amplicon generated from the small RNA fraction matched the specific melting profile of the miR-686 positive control (Figure 2), indicating that specific amplification was achieved only with the small RNA fraction.

**Figure 4.**
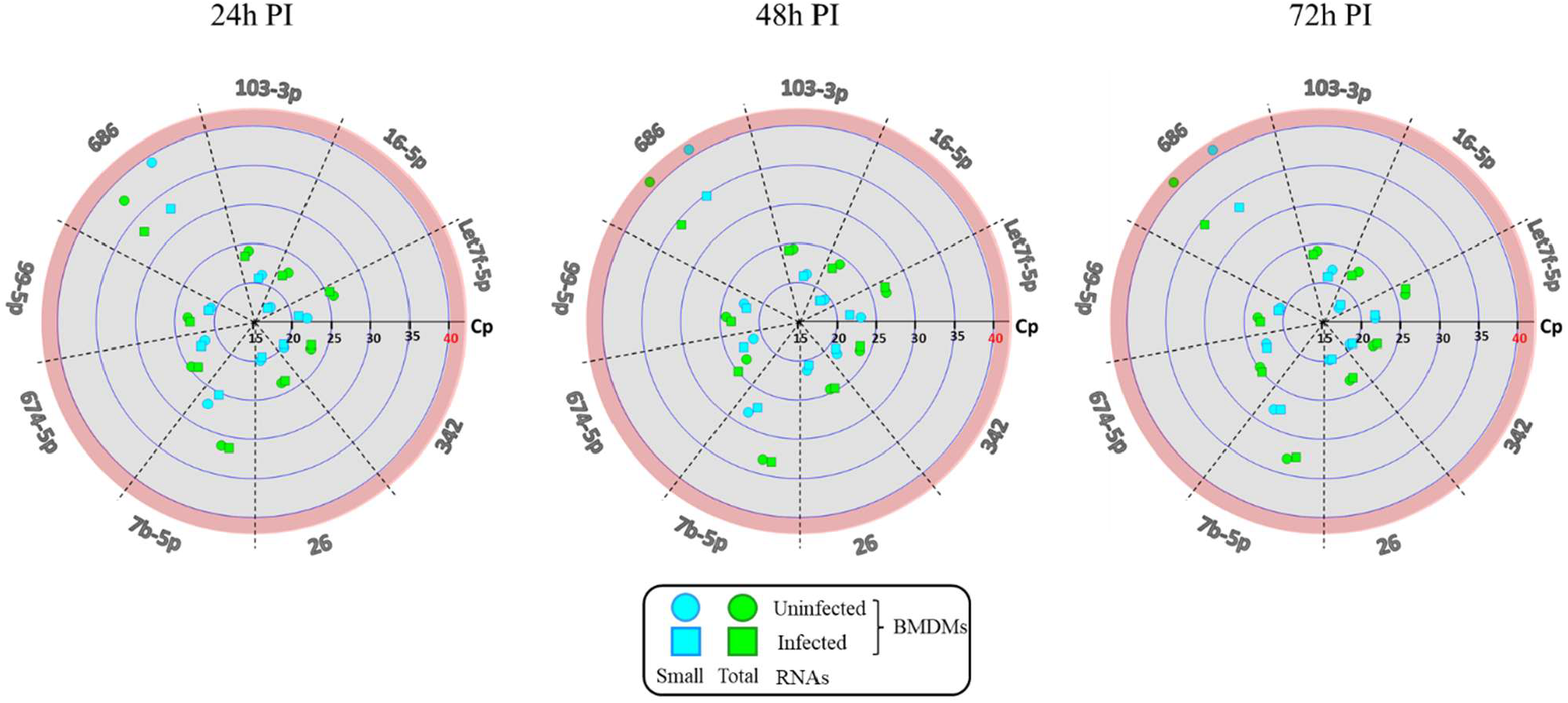
Comparison of Cq values obtained with small and total RNA fractions isolated from *L. amazonensis* infected and uninfected BMDMs at 24h, 48h and 72h post infection. Small RNA fractions consistently yielded lower Cq values than total RNA fractions in RT-qPCR, with the exception of miRNA-686, for which Cq values were comparable between the two RNA fractions.

## Conclusions

The use of small RNA fractions yields lower Cq values across all target miRNAs and experimental conditions, followed by total RNA fraction, while the large RNA fraction proves to be the least suitable for this type of analysis both in terms of Cq values (Figure S1) and melting curve specificity (Figure S2). For low-expressed miRNAs, such as miR-686 in UI samples and miR-7b-5p in our experimental model, specific amplicons can be detected using small RNA fractions applying Protocol 2 as specified in the methods section. These findings highlight the critical importance of RNA fraction selection and protocol optimization in RT-qPCR assays targeting low-abundance miRNAs. Specifically, the use of small RNA combined with the modified Protocol 2 emerges as the preferred approach to maximize both sensitivity and specificity. MiRNAs represent promising diagnostic and prognostic biomarkers, as well as therapeutic targets, particularly in infectious diseases like leishmaniasis, where the parasite employs complex strategies to alter host miRNA expression. The optimization approach presented in this study is particularly valuable for detecting biologically meaningful changes in weakly expressed miRNA targets. Further studies will aim to validate and extend these findings across broader experimental contexts.

## Acknowledgment

We acknowledge the financial support of the Institut Pasteur (Paris) and the Institut national de la santé et de la recherche médicale (INSERM). Sara Maestrini is supported by the Department of Biomolecular Sciences, University of Urbino Carlo Bo. The funders had no role in study design, data collection and analysis, decision to publish, or preparation of the manuscript.

