## Supplementary figures for "Enhanced detection of low-expressed miRNAs in *Leishmania*-infected macrophages through RNA fractionation and RT-qPCR optimization"

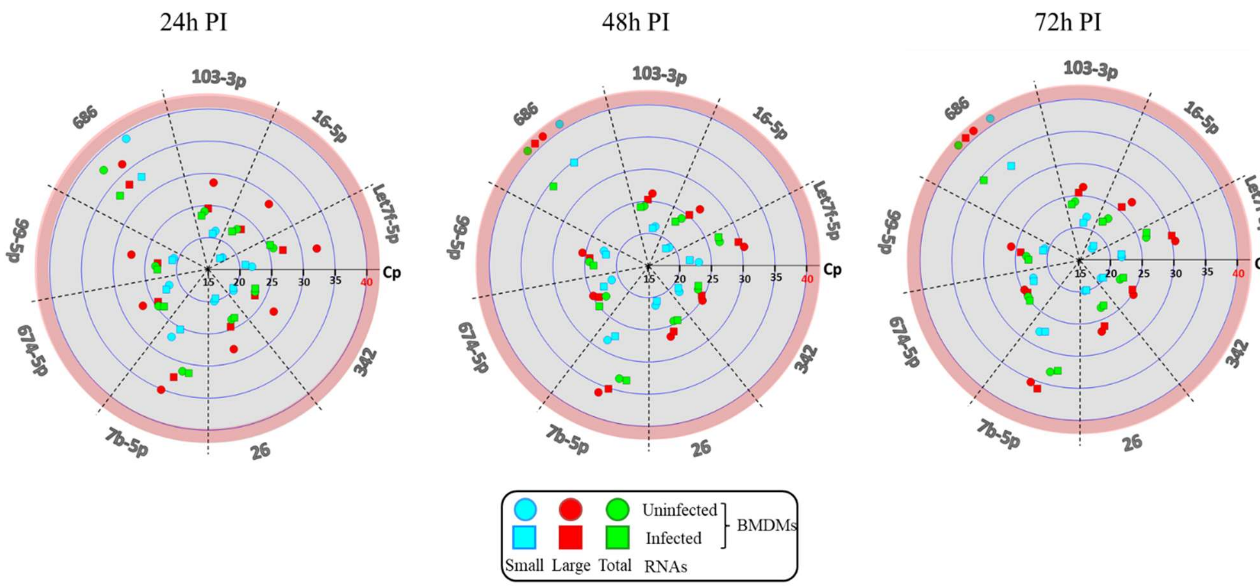


**Figure S1: Comparison of Cq values obtained with small, large and total RNA fractions in *L. amazonensis* infected and uninfected and BMDMs samples 24h, 48h and 72h post infection.** Small RNAs as RT-qPCR input gave lower Cq values than total and large RNA, except for miRNA-686, for which Cq values of small and total RNAs are comparable.

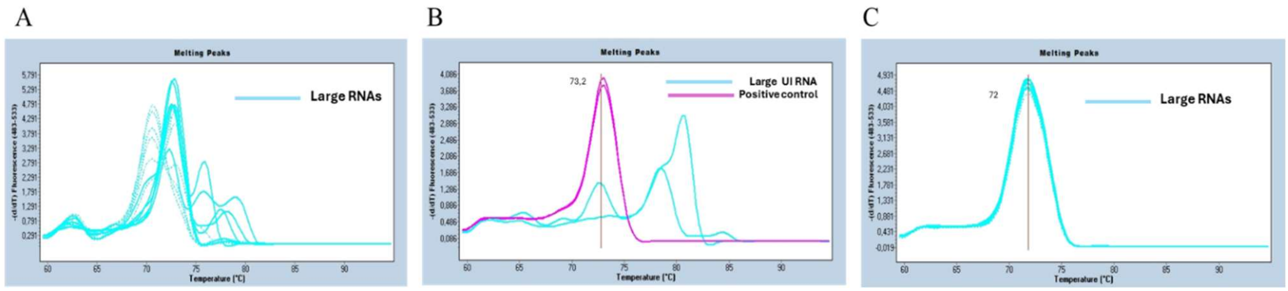


**Figure S2: Melting curves obtained with large RNA fractions in *L. amazonensis* infected and uninfected BMDMs samples for highly and low expressed miRNA targets.** A) melting curves of miR 7b-5p in large uninfected (UI) BMDMs and infected (I) BMDMs 24h, 48h and 72h PI; B) melting curves of miR-686 in large RNA of UI BMDMs 72h PI; C) melting curves of miR-26b in large UI BMDMs and I BMDMs 24h, 48h and 72h PI. The large RNA fraction is unsuitable for miRNA RT-qPCR due to non-specific products for low-expressed targets.
